# Net conversion calculations of catabolic pathways

**DOI:** 10.64898/2026.09.04.749342

**Authors:** Frank Bruggeman, Maaike Remeijer, Christoff Odendaal, Rebeca Gonzalez-Cabaleiro

**Affiliations:** Systems Biology Lab, A-LIFE, AIMMS, Vrije Universiteit, Amsterdam; Environmental Biotechnology Section, Department of Biotechnology, Delft University of Technology, the Netherlands

## Abstract

**Motivation:** The stoichiometry (or net conversion) of a catabolic pathway is an often used principle of biochemistry. It expresses the molar yield of charged energy carriers (e.g. ATP) and catabolic products (e.g. lactate) on the energy source (e.g. glucose). Product yields are engineering targets of metabolic engineering and used in microbial ecology to assess energy metabolisms of microbial species. For a single species under a single condition, the catabolic pathway is frequently assumed fixed, while there might be multiple options encoded in its genome. To find these options, the manual (heuristic) methods that have been used for decades fall short.

**Results:** In this paper, we explain how net conversions can be calculated from reaction stoichiometries of a metabolic network and evaluated using thermodynamic information. We start with an (old) heuristic method. Next, we explain we relate the net conversions of metabolic networks to their elementary flux modes (EFMs) and show that a single EFM gives rise to a single net conversion. EFMs are mathematical objects that are computable with existing software. We use these to illustrate how all net conversions of complex (pan-)metabolic networks can be computed. We consider examples from aerobic and anaerobic microbiology. Then, we introduce the parameter Ω, the driving force per unit flux, which allows for the thermodynamic comparison of pathways. To calculate Ω, only the standard Gibbs free energy potential, i.e., Δ*G^m′^* of the net conversion and its corresponding EFM are required. A generic workflow (and all underlying Python code) are provided, as well as applications to perform the workflow without coding. We also provide those software packages that automate our methods.

**Impact:** This paper serves as an illustration of how modern computational systems biology can be used to automate the calculation of net conversions in microbial ecology and metabolic engineering. We hope that this paper inspires future metabolism research using quantitative, rigorous methods.

## Introduction

The main function of catabolism is to provide anabolism with charged energy carriers such as NAD(P)H and ATP [1]. This is accompanied by the formation of catabolic byproducts. These catabolic pathways can each be expressed as a net conversion, specifying the yield of charged energy carriers and catabolic products on glucose [1, 2].

Many different catabolisms have been discovered in nature, for example the production of lactate from glucose [3] (glucose + 2 ADP + 2 Pi → 2 lactate + 2 ATP + 2 H_2_O), the production of propionate and acetate from lactate [4] (3 lactate + 2.5 ADP + 2.5 Pi → acetate + 2 propionate + CO_2_ + 3.5 H_2_O + 2.5 ATP), or the conversion of carbon dioxide and hydrogen to methane by methanogens [5] (4 H_2_ +CO_2_ + 1.5 ADP + 1.5 Pi → CH_4_ + 3.5 H_2_O + 1.5 ATP).

Similar conversions might also be carried out via different routes. For example, certain species may rely on the Embden-Meyerhof-Parnas (EMP) pathway to convert glucose into ethanol and carbon dioxide to synthesize ATP (glucose + 2 ADP + 2 Pi → 2 ethanol + 2 CO_2_ + 2 ATP + 2 H_2_O) while others use the Entner-Doudoroff (ED) pathway instead, generating 1 fewer ATP per glucose (glucose + ADP + Pi → 2 ethanol + 2 CO_2_ + ATP + H_2_O).

Microbes adapt their catabolism to environmental conditions by shifting their net conversions. *Escherichia coli* (*E. coli* ) can, for instance, carry out several glucose catabolisms [1]. Aerobically, it can either fully respire glucose into carbon dioxide, break it down only partially into acetate, or mix the two strategies. Anaerobically, *E. coli* synthesizes ATP from glucose via mixed-acid fermentation with ethanol, acetate and formate as catabolic products. Understanding why certain catabolisms are preferred by microbes and under which conditions is important to predict the metabolic strategies of microbes. This has been a central objective in microbial physiology and ecology for decades [2].

Net conversions are a property of a metabolic network at steady-state [6]. This is the state of the metabolism during balanced growth of (microbial) cells [7]. The network then displays time-invariant rates of substrate uptake and product formation such that the synthesis and consumption rates of each pathway intermediate are balanced. As a result, pathway intermediates do not accumulate or deplete over time. The requirement of balancing the synthesis and consumption rates of redox equivalents such as NADH is of particular importance in anaerobes, because these need to be recycled back to the oxidized equivalent, in this case NAD^+^. In the absence of a high potential external electron acceptor such as oxygen, pathway intermediates are used as electron acceptors; in these situations, redox balancing becomes a powerful constraint on the variety of possible conversions [8]. A recent textbook reviews several common anaerobic catabolic conversions that have been mapped onto underlying pathways that maintain redox balance [2].

Steady-state net conversions can be determined from the stoichiometry of the reactions in the network [6]. Manual methods rely on linear algebra or direct balancing of all the synthesis and consumption rates of pathway intermediates, particularly of redox equivalents. While some authors make use of genomic or gene expression data to identify probable underlying pathways [9], often common pathways are simply assumed based on measured product profiles [10].

Net conversions can also be identified from reaction networks computationally using elementary flux modes (EFMs) [6]. An EFM is a mathematical concept of the set of feasible steady-state rate values of a metabolic network and provides an unambiguous definition of what constitutes a metabolic pathway [11]. Their mathematical definition and properties can be found in Gagneur & Klammt [12]. Python packages exist for their calculation given the stoichiometric information of the pathway’s reactions, e.g. EFMTool [13].

An EFM corresponds to a set, or vector, of reaction rate values at steady state (called fluxes). None of the reactions associated with this set are redundant (so the network is minimal, no reaction can be removed without violating the steady state condition) and EFMs therefore have a single degree of freedom, i.e. all flux values can be determined if one is known. These reactions form a minimal metabolic network, the ‘EFM network’. Only one net conversion is associated with a given EFM network. Different EFM networks can however give rise to the same net conversion. A given catabolic network may be capable of catalyzing many net conversions, each associated with one or more EFM networks.

In this work, we show the net conversions studied by microbial ecologists (e.g. [2]) and biotechnologists coincide with the principles of EFMs, because the set of EFMs of a metabolic network can be recombined to yield all possible conversions. In fact, while not a prerequisite for our argument, it is interesting to note that authors have previously shown that single EFMs, instead of mixed strategies, always represent the most resource-efficient and highest yield (e.g., of ATP) solutions of a network [14, 15]. EFMs therefore provide a way of mapping conversions onto feasible networks in an unbiased way, potentially allowing for the discovery of previously overlooked pathway stoichiometries in microbial ecology. Computation of EFMs therefore offers a useful computational method for microbial ecologists and microbiologists alike.

This paper is organized as follows. We start by explaining two versions of the manual method, using glucose fermentation to lactate as an example. Next, we introduce EFMs and show several examples of them together with their unique net conversions. We illustrate how those were determined from the reaction stoichiometries of the pathways using a Python script (using EFMtool [13]) and a software tool which we developed. Then we show how the Gibbs free energy change (Δ*G*) of a net conversion can be calculated by hand or by using a Python script (by means of eQuilibrator [16]). We introduce a parameter, Ω, which we use to compare different metabolic networks. Finally, we show how all the elementary net conversions of a complex (pan-) metabolic network can be calculated. This is an application where the manual method becomes too laborious. Throughout this paper, we use examples that are relevant both for microbial ecology and metabolic engineering.

## Results

### Steady-state catabolic networks, external and internal metabolites

Net conversions are properties of metabolic networks operating under steadystate conditions. A steady state requires that the concentrations of the substrates and products of the pathway are kept fixed. For example, for glycolysis shown in Figure 1, this implies that glucose, ATP, ADP, Pi and lactate have constant concentrations. Such metabolites are defined as “external” metabolites: they are either net-consumed or -produced by the network. All the remaining reactants are “internal” and function as pathway intermediates. They are synthesized and consumed at equal rates, such that their concentrations remain constant in time. When we focus on ATP as the sole charged energy product of catabolism. The synthesis and consumption of all other electron carriers, such as NAD(P)H, are therefore balanced. This is generally the approach taken in the literature (e.g., [2]) and in this paper. Because the external metabolites are net substrates and products of the pathway, only they appear as reactants in the net conversion reaction.

**Figure 1.**
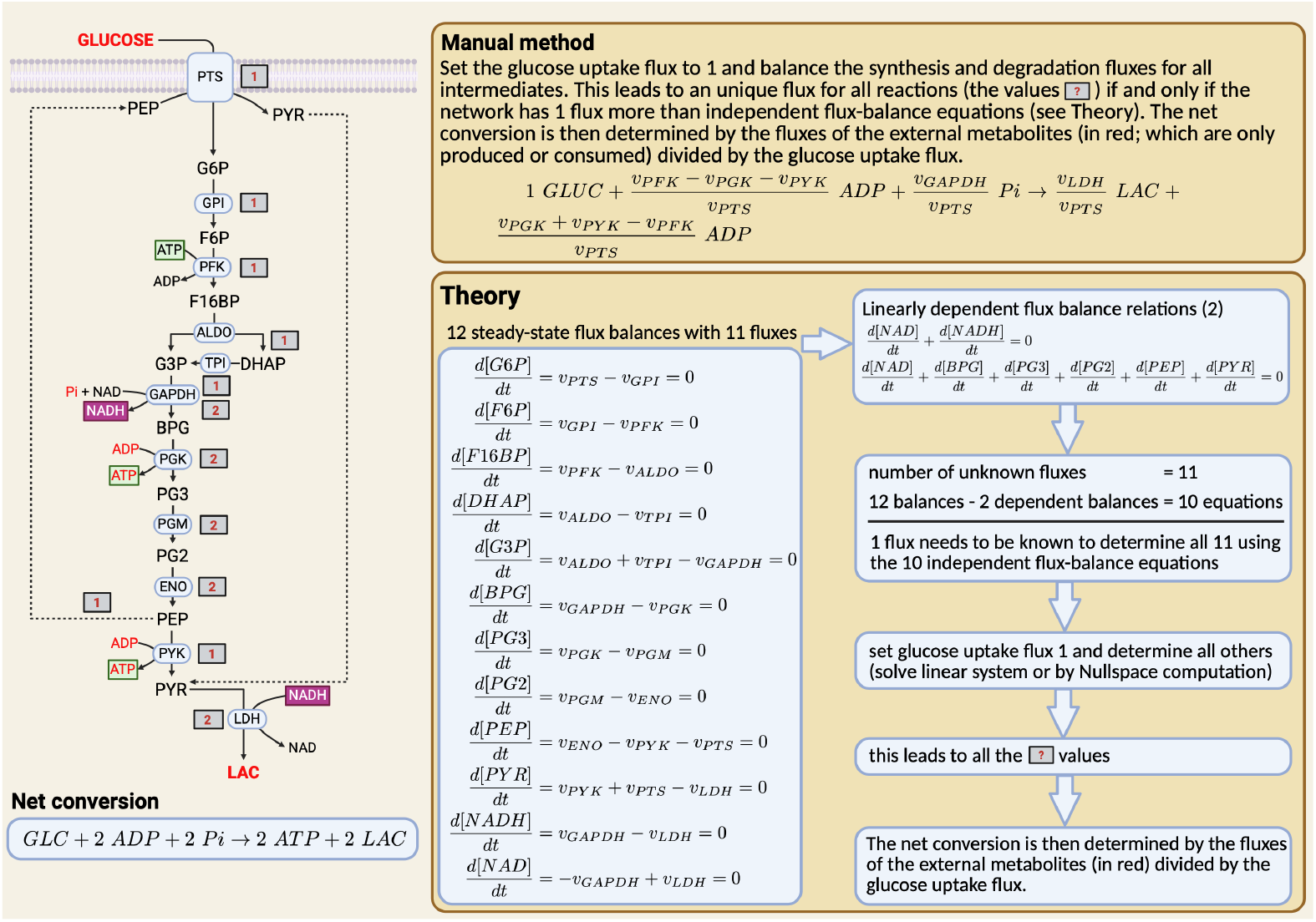
Illustration of the manual method and underlying theory for determining the net conversion of a catabolic pathway. The Embden-Meyerhoff-Parnas pathway is shown with a bacterial phosphotranseferase system as the glucose import mechanism. The shown pathway was deliberately chosen to have only one independent flux. This conversion is, for instance, catalyzed by the bacterium *Lactococcus cremoris* (*L. cremoris* ) under aerobic conditions and by some *E. coli* strains under anaerobic conditions. This network has 5 external metabolites (glucose, Pi, ADP, ATP and lactate), 12 internal metabolites (G6P, F6P, F16BP, G3P, DHAP, BPG, PG3, PG2, PEP, PYR, NADH, and NAD), and 11 enzyme-catalysed reactions (PTS, GPI, PFK, ALDO, TPI, GAPDH, PGK, PGM, ENO, PYK, LDH). A representative unit for all the flux values are 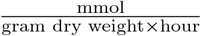. Water molecules and protons are not considered in the reaction equations.

### Stepwise methods: heuristic, linear-algebra based and computational

We will consider two manual methods for determining the net conversion of a steady state catabolic pathway: the ‘heuristic method’, involving manual flux balancing, and the ‘mathematical method’, involving linear-algebra-based flux balancing.

#### Heuristic method: manual flux balancing

The heuristic method starts from the network diagram of the catabolic pathway. It determines the flux values of all the reactions by balancing (equalizing) the synthesis and degradation fluxes of all internal metabolites, after fixing one of the flux values at a reference value of 1. Typically, the uptake rate of the energy source (external metabolite; glucose in Figure 1) is set to 1. Depending on the network, it may turn out that more than one set of flux values leads to a steady state. In that case, the catabolic pathway considered has more than one EFM, and might have multiple possible net conversions. That is not the case for the pathway shown in Figure 1: knowing one flux value suffices for determining the values of all others uniquely.

Thus, we start by assigning the rate of the glucose uptake reaction (the phosphotransferase system, or PTS, in this case), denoted by *υ*_*P T S*_, a value of 1 (mmol*/*(gram dry weight *×* hour), if it is a specific flux). The reader will appreciate that each subsequent reaction has only one possible rate value, as each intermediate must be produced and consumed at the same rate (Figure 1). The uniqueness of NADH is also apparent, as it is not consumed and produced by immediately adjacent reactions, but produced by GAPDH and consumed further downstream by LDH. In this way, the carbon conversions downstream of GADPH have to run at a rate that ensures that all NADH produced can be reoxidized to NAD^+^. A step-by-step analysis of this system is described in Appendix S1.

When all the steady-state flux values have been determined (i.e., the values in the grey boxes in Figure 1), we can determine the stoichiometry of the net conversion. This requires only the flux values of the external metabolites: GLC, ATP, ADP, Pi, and LAC. We start by calculating the ratio of their net consumption or production flux over the glucose uptake flux. The net fluxes are

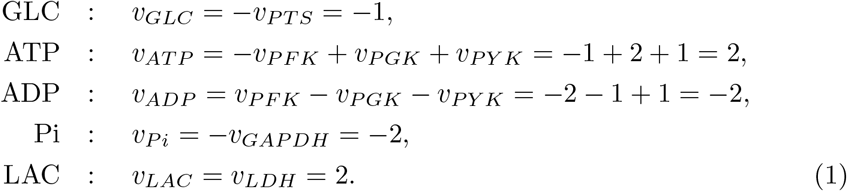

Thus, GLC, ADP, and Pi are consumed by the network and occur on the left hand side of the net conversion, whereas ATP and LAC are produced and occur on the right hand side. The net conversion per mole of glucose equals,

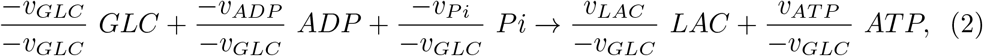

and therefore,

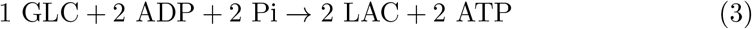

Thus, the molar yield of lactate and ATP on glucose is 2.

#### Mathematical method: linear-algebra-based flux balancing

The mathematical method for determining the net conversion requires that the network has a single independent flux. This implies that all the fluxes in the network have a unique flux value at steady state when the uptake flux value of the energy substrate has been set to 1. This applies to the metabolic pathway shown in Figure 1.

Confirming that this network has only a single independent flux is best done by analyzing the stoichiometric matrix **S** of the catabolic pathway. This matrix contains the stoichiometry of all the reactions in the network. (For instance, the stoichiometry of the glucose import reaction (by PTS) of the network is *GLC* + *PEP* → *G*6*P* + *PY R*.) The number of rows of matrix **S** equals the number of internal metabolites, its number of columns equals the number of reactions, and its multiplication with the vector of the reaction rates equals the vector of all the rates of change of the concentration of the internal metabolites:

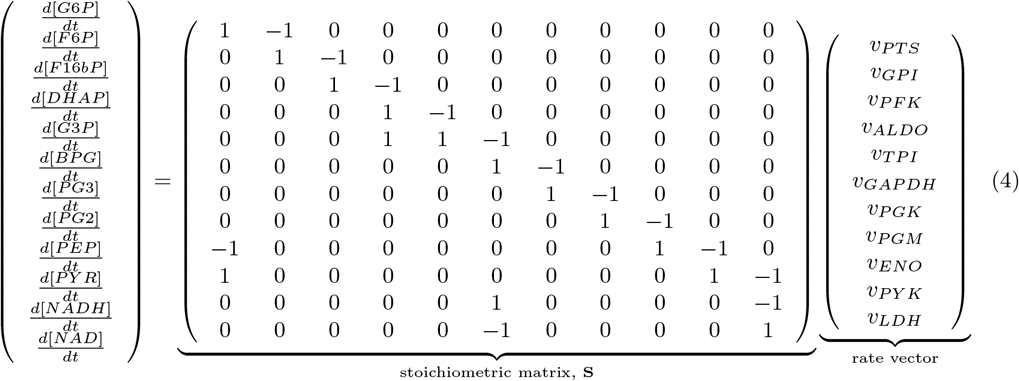

The value of the change of each metabolite over time equals zero at steady state. Therefore, at steady state, this equation equals a vector of zeros, and the flux vector contains the steady state rate values. Supplementary file S1 shows this calculation in a Python notebook.

A metabolic network has a single independent flux when the number of reactions in the network (11 in Figure 1) minus the rank of the stochiometric matrix **S** equals 1. The rank of the stoichiometric matrix indicates the number of linearly independent rows of the stoichiometric matrix and equals 10 in this example.

An aspect not required for the computation of the net conversion from the stoichiometric matrix is that its linearly dependent rows indicate the constancy of sums of concentrations of internal metabolites [17]. Matrix **S** has 12 rows and its rank equals 10, so 2 rows are linearly dependent. These can be retrieved from a linear combination of the 10 linearly independent rows. One such linear combination is that *d*[*NADH*]*/dt* = −*d*[*NAD*]*/dt* which implies that *d*([*NADH*] + [*NAD*])*/dt* = 0 and, hence, the sum of the concentrations of *NADH* and *NAD* remain constant. The other is shown in Figure 1. In the glycolysis of Figure 1, the linear combinations of the flux balance equations lead to the following constant pools of metabolites,

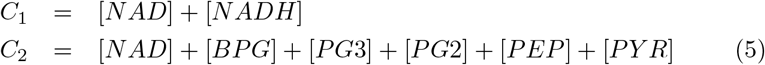

Now that it is established that the network shown in Figure 1 has only a single independent flux value, the rate vector at steady state can be determined by calculating the nullspace of the stochiometric matrix. (The nullspace now equals a vector and not a matrix because the number of its columns equals the number of independent fluxes.) The nullspace vector is unique up to multiplication; so multiplying all its entries with the same constant preserves steady state. Hence, division of all its entries by the value of the import rate of the energy substrate (to normalize its value to 1) is again a steady-state rate vector. Now that we know the steady-state rate vector we can proceed to determine the net conversion as explained at the end of the last section (Equation 1).

#### Computational method: Python and app-based calculation

When the considered metabolic network is large, and contains branches and cycles, both the manual and linear-algebra based methods become quickly complicated and laborious. EFMTool can find all EFMs in a stoichiometric matrix (up to a reasonable size). We developed a small pipeline to convert an intuitive spreadsheet into a stoichiometric matrix and subsequently calculate all EFMs. The steps required are described in the Methods. This pipeline is available as an interactive Python notebook (Supplementary file S3), which uses a spreadsheet defining the reactions and metabolites as input (Supplementary file S2). Additionally, we present an application, the NetConverterApp, that uses the same steps. Because the generation of the spreadsheet is cumbersome, we also present an app, the KEGGBuilderApp, that takes KEGG reaction or module identifiers [18] to generate the spreadsheet.

### Determination of the Gibbs free energy change of a net conversion

The net conversions of (heterotrophic) catabolic networks can generally be written as the sum of two elementally balanced reactions: the conversion of the carbon source into the catabolic product and the phosphorylation of ADP. For instance, the conversion of glucose into formate, acetate and ethanol yielding 3 ATP per glucose can be written as the following sum of reactions,

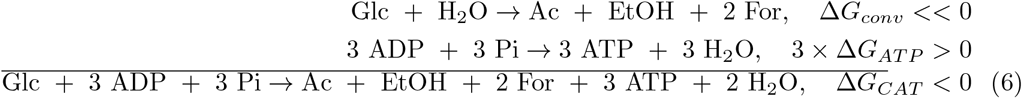

A characteristic of such net conversions is that the breakdown of the carbon source (glucose in this case) liberates Gibbs free energy (i.e., Δ*G*_*conv*_ *<<* 0 and 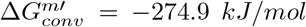; calculated with eQuilibrator’s web interface [16]). This free energy is (partially) used to phosphorylate ADP to yield ATP, a reaction which demands Gibbs free energy to proceed (i.e., Δ*G*_*AT P*_ *>* 0 and 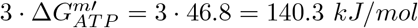). The net conversion can be written as a sum of these two elementally balanced reactions and has a negative free energy (i.e., Δ*G*_*CAT*_ = Δ*G*_*conv*_ − 3 · Δ*G*_*AT P*_ *<* 0 and 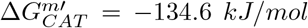). The values of Δ*G*_*conv*_ and Δ*G*_*AT P*_ were determined from the formation Gibbs free energies of the reactants, which are dependent on reaction conditions (now standard conditions), and their concentrations. The Gibbs free energy change of any reaction or net conversion can be calculated with eQuilibrator [16], using its web interface or its Python API, which we use in the Python notebooks and the apps we provide as Supplementary Information. A more general introduction to thermodynamics is given in Appendix S2.

Thus an amount of Δ*G*_*CAT*_ Gibbs free energy is dissipated and lost as heat. This waste has a benefit; the rate of a net conversion (per unit invested protein) is higher when its Δ*G* is more negative [19]. (This rate is zero when Δ*G* = 0, at thermodynamic equilibrium.) Therefore, a lower expenditure of Δ*G*_*conv*_ on ATP synthesis would therefore increase the rate of the net conversion. This effect may in some cases explain a trade off between rate and yield of catabolic pathways.

The relationship between the Gibbs free energy change of the net conversion (Δ*G*_*cat*_) and the Gibbs free energy changes of its underlying pathway reactions (Δ*G*_*i*_) is given by (Figure 2):

**Figure 2.**
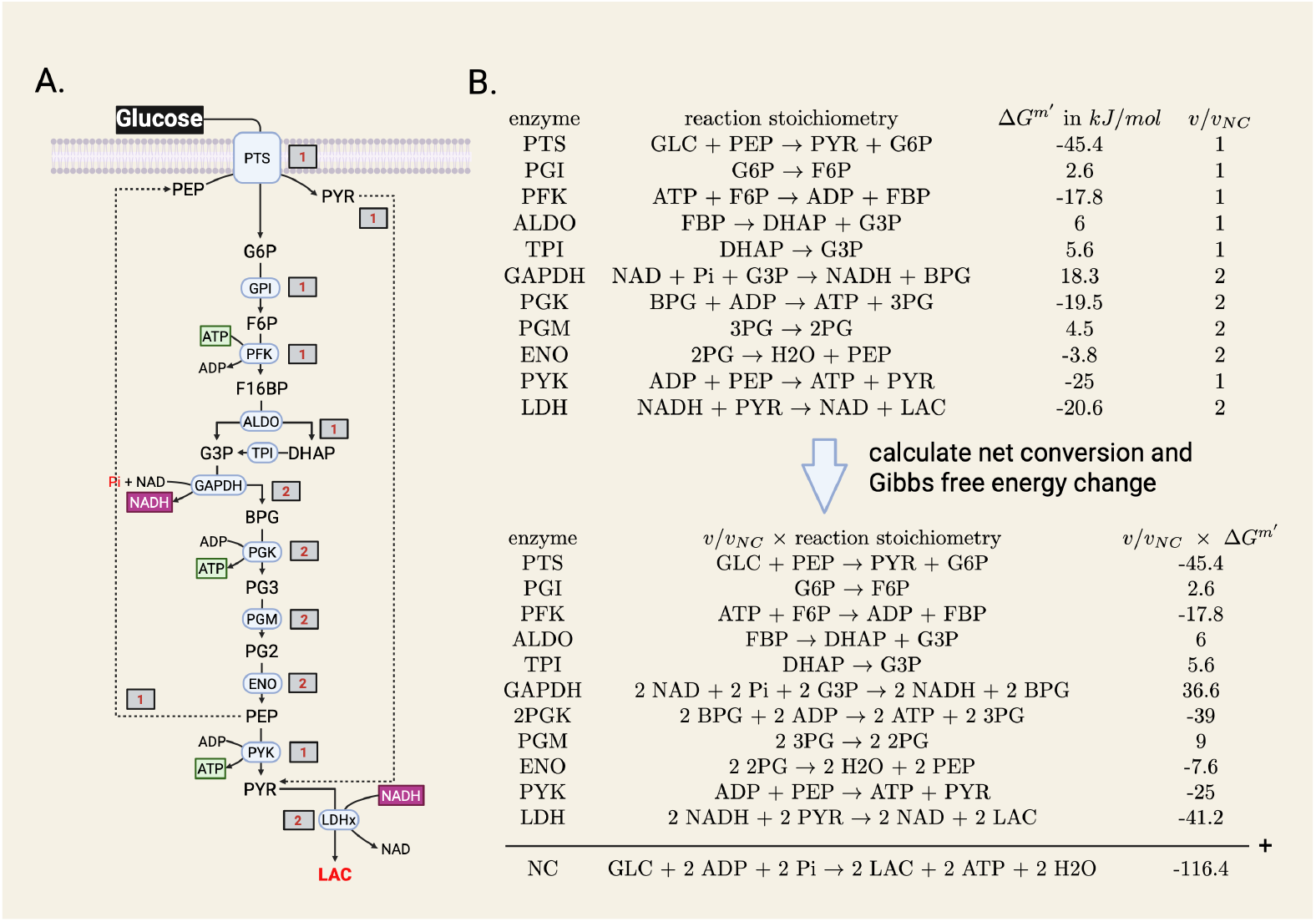
Expressing the net conversion and its standard Gibbs free energy change Δ*G*^0*′*^ or Δ*G*^*m′*^ in terms of the underlying reaction properties. This figure shows: i. how the net conversion can be obtained by summing all the reaction stoichiometries multiplied by their normalised rate value, and ii. how the standard Gibbs free energy change is obtained by summing all the standard Gibbs free energy changes of the reactions multiplied by their normalised rate value.

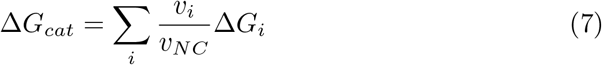

with *v*_*i*_ and Δ*G*_*i*_ as the steady-state rate and Gibbs free energy change of reaction *i* and the *v*_*NC*_ as the steady-state rate of the net conversion (equal to 1 when the influx of the energy source is set to 1). In words, the Δ*G* of the net conversion is equal to the fractional-flux weighted sum of the Δ*G* of the individual reactions. In Figure 2, we illustrate this equation for the calculation of the 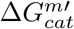 calculation for the example network, which was also analysed in Figure 1. The 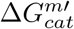 is the sum of 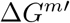 values of the individual reaction steps, weighted by their normalised flux value.

To analyze the thermodynamic bottlenecks of pathways, the max-min driving force (MDF) is often calculated [20]. In this approach, the minimum driving force (-Δ*G*^*m′*^) across all reactions is maximized in a linear program, given the individual Δ*G*^*m′*^ values of reactions and upper and lower bounds for the internal metabolite concentrations. This approach has two limitations: the Δ*G*^*m′*^ values of individual reactions can carry substantial uncertainty, for example, when ferredoxin or other complex electron carriers or transport processes are involved, or when intracellular metabolite concentrations are poorly characterized, particularly for non-model organisms.

We introduce a new thermodynamic property of a pathway: the Δ*G*^*m′*^ per unit flux (Ω), defined as follows:

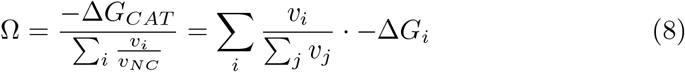

The derivation can be found in Appendix S3. To calculate Ω, no knowledge of Δ*G*^*m′*^ of individual reactions or of intracellular concentrations is required. Ω captures the free energy dissipated per reaction event (−*v*_*i*_Δ*G*_*i*_), weighted by the flux sum ( ∑_*i*_ *v*_*i*_). The value of Ω is equal to the value of the MDF result when no concentration bounds are used for the MDF computation, and the upper bound for any MDF computation. We show this in the appendix.

Intuitively, for a linear pathway with equal fluxes, Ω = −Δ*G*_CAT_*/n*_*R*_, which represents the upper bound on Ω. When reactions carry unequal fluxes, the MDF solution distributes the total free energy dissipation proportionally, and Ω corresponds to the driving force at the bottleneck reaction: the one carrying the lowest flux (normalised to 1). We show this for a few examples in Appendix S4. In addition, Ω can be used to rank the thermodynamic properties of the EFMs (Figure S1).

### EFMs can be complicated pathways

In Figure 3 and 4, we show eight catabolic pathways of a range of microbial species. Each pathway has a single independent flux. Their net conversions can be obtained with the nullspace, manual method, the Python pipeline (Supplementary file S4 and S5), or the app. The values for Δ*G*^*m′*^ are also shown and were computed with eQuilibrator.

**Figure 3.**
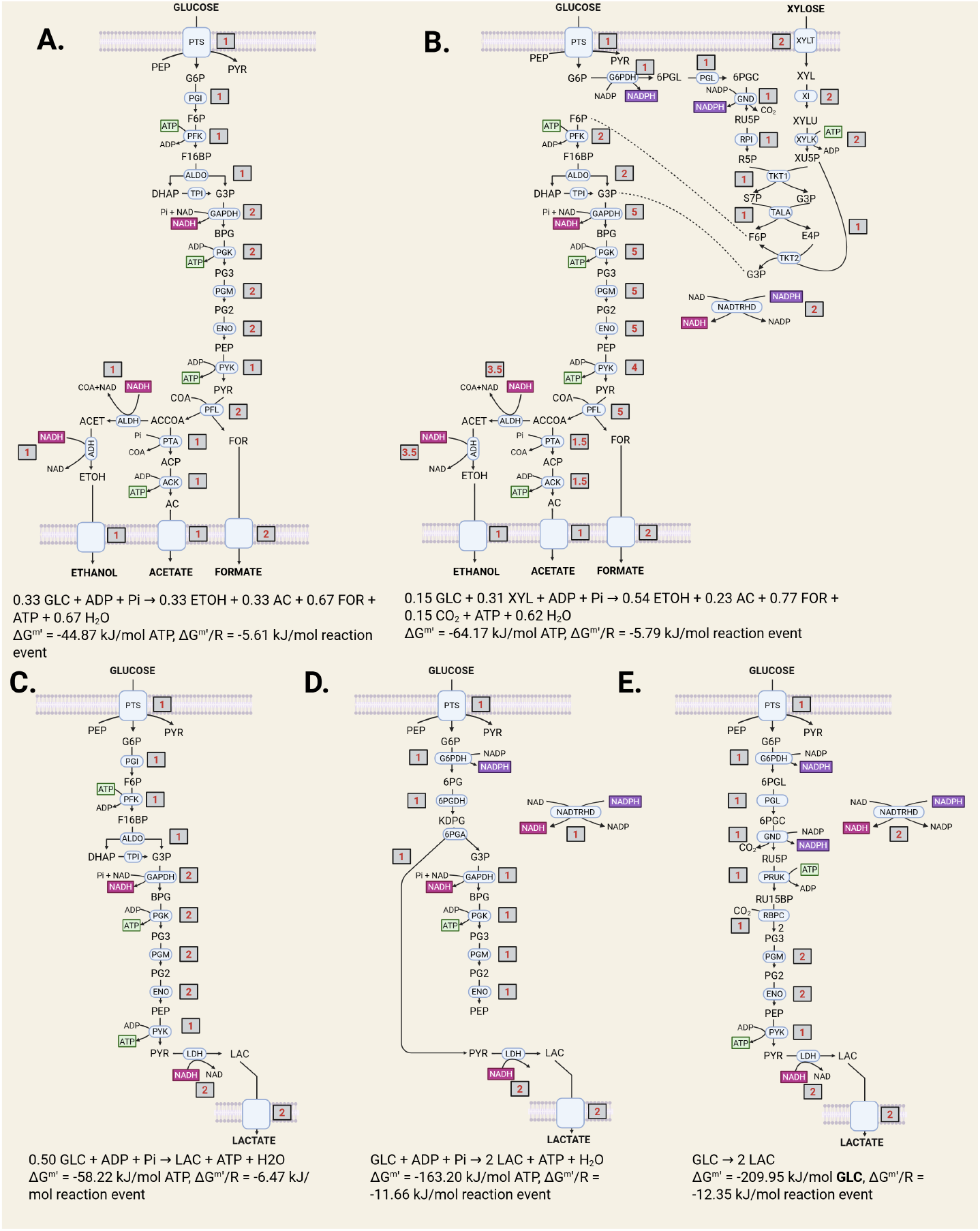
Five examples of metabolic networks involving glycolysis with a single independent flux, each containing at least 1 irreversible reaction, and therefore all elementary flux mode networks. Five metabolic networks are shown together with their net conversion and associated 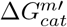. They each have a single independent flux and can run only in one direction due to the occurrence of at least one irreversible reaction, which ensures that they are all elementary flux modes. The values in the grey boxes indicate the steady-state flux values. The overall equations are normalized for the production of 1 ATP. From this equation, the 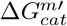 was additionally calculated. We provide all the Python scripts underlying these computations. (A) Mixed acid fermentation with the Embden-Meyerhof-Parnas (EMP) glycolysis. (B) Designed network for coconsumption of glucose and xylose, involving EMP glycolysis and the pentose phosphate pathway. (C) Lactic acid production with the EMP glycolysis. (D) Lactate production with the Entner-Doudoroff (ED) glycolysis. (E). ATP-neutral lactate production that utilizes the ‘Calvin shunt’, i.e. some enzymes from the Calvin cycle.

**Figure 4.**
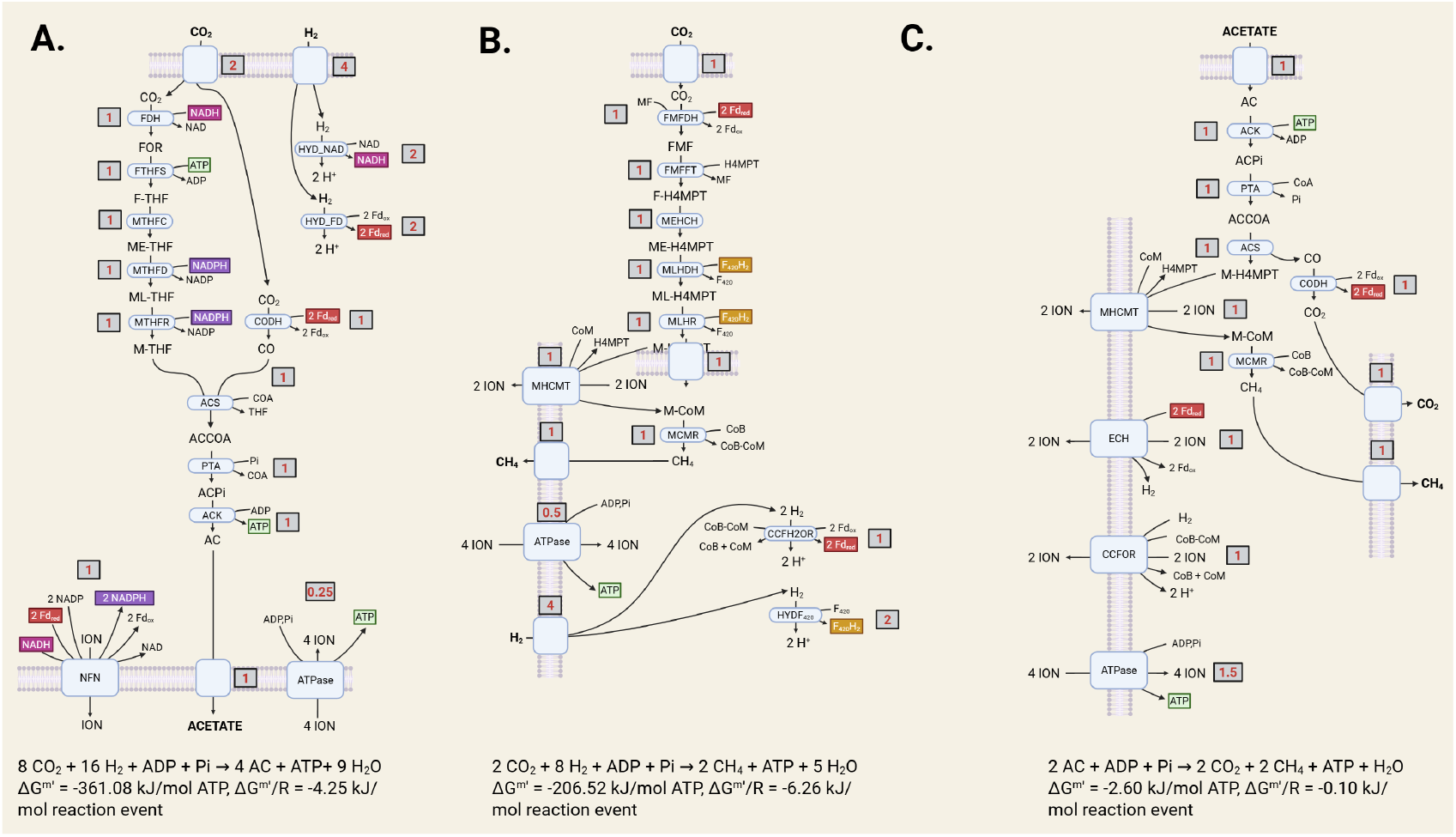
Three examples of non-glycolytic metabolic networks with a single independent flux, each containing at least 1 irreversible reaction, and therefore all elementary flux mode networks. Three metabolic networks are shown together with their net conversion and associated 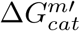. They each have a single independent flux and can run only in one direction due to the occurrence of at least one irreversible reaction, which ensures that they are all elementary flux modes. The values in the grey boxes indicate the steadystate flux values. The overall equations are normalized for the production of 1 ATP. From this equation, the 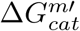 was additionally calculated. We provide all the Python scripts underlying these computations. (A) Wood-Ljungdahl pathway. (B) Hydrogenotrophic methanogenesis. (C). Acetoclastic methanogenesis.

These examples indicate that pathways with a single independent flux can be quite complicated. A computational method that can compute the net conversions of a catabolic network, regardless of its number of independent fluxes, is therefore very useful. We encourage the reader to apply the methods described above to the networks drawn in Figure 3 and 4. All networks considered so far (Figure 1, 3 and 4) are EFMs. In this work, we demonstrate the method for only relatively simple EFMs. In genome scale metabolic networks, EFMs that include growth typically have a few hundred reactions.

### Combinatorial explosion of the number of EFMs

A single network can consist of many EFMs due to a ‘combinatorial explosion’. An example of such a combinatorial explosion case is shown in Figure 5A. It shows variations of glycolysis, that give rise to alternative routes: glucose can be imported via two different import reactions, phosphorylated and split via EMP or ED, the resulting G3P can be converted via GAPDH or GAPN, and finally three different fermentation routes are considered from pyruvate. This leads to 2×2×2×3=24 alternative EFMs (Figure 5B). Their net conversions and properties are shown in Figure 5. These calculations were done with the provided Python notebook and computational tool, which starts from the stoichiometric description of each of the involved reactions (Supplementary file S6 and S7).

**Figure 5.**
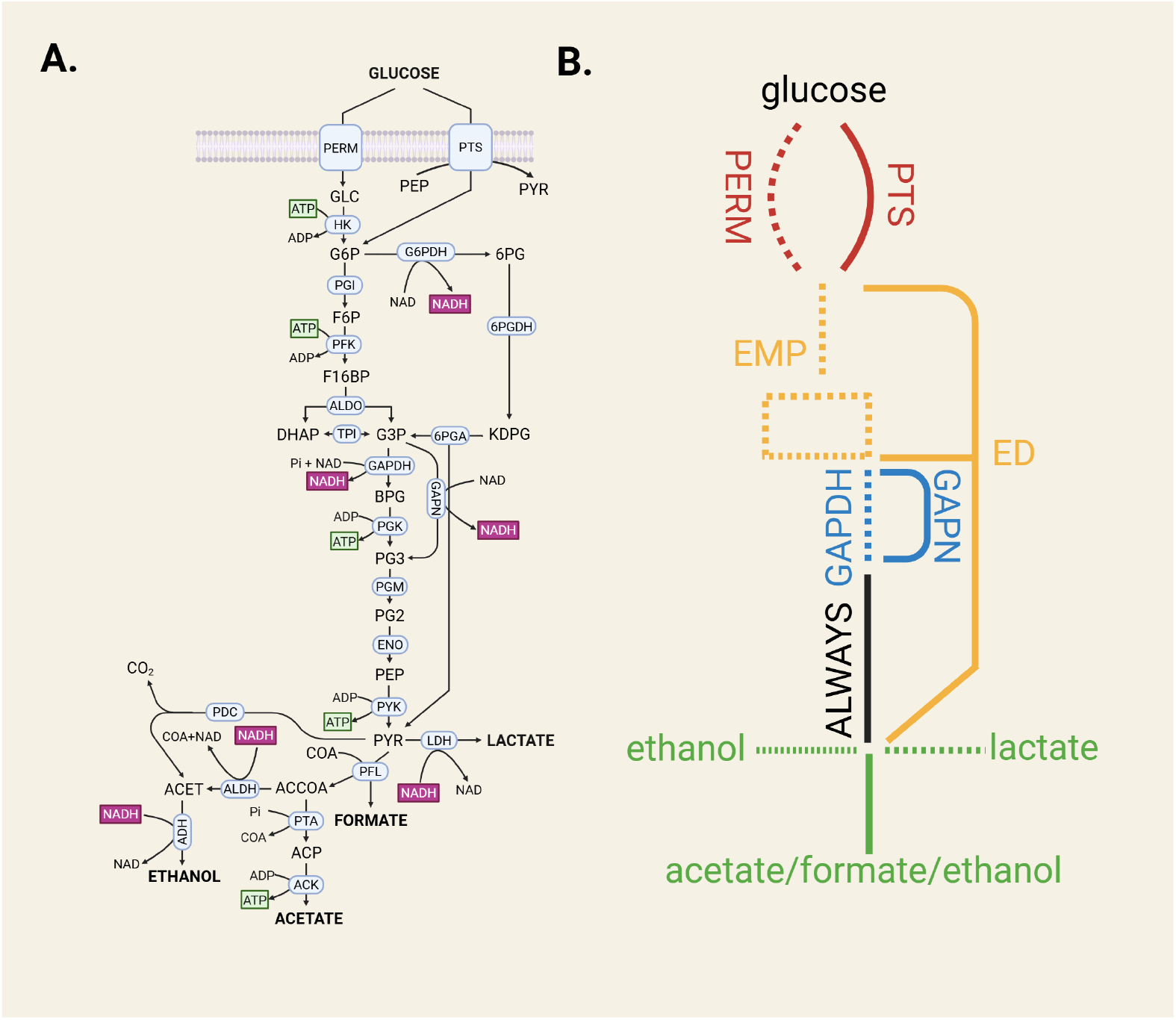
A glycolytic network to show the combinatorial explosion of EFM enumeration. (A) All reactions in the metabolic network and a schematic overview of the combinatorics of the network. Two options for glucose transport are available (permease and the PTS system), two options for the conversion of G6P into G3P (EMP and ED), two options for G3P to 3PG (GAPDH + PGK and GAPN) and three product pathways (ethanol, mixed acid fermentation and lactate). This yields 2 *×* 2 *×* 2 *×* 3 = 24 EFMs.

The combinatorial explosion of the network shown in Figure 5A was deliberately kept small for illustrative purposes. Realistic metabolic networks of intermediate size with many bypasses and branches can easily lead to thousands of solutions. For example, if the reactions from the EFM examples in Figure 3 and 4 are combined, the network contains 3183 EFMs, as can be seen from one of the pre-loaded examples of the net conversion app. Another example is the ‘ecolicore’ network, which contains 95 reactions, has approximately 272 million EFMs [21]. At the genome scale, enumeration of EFMs remains computationally infeasible. Many of the EFMs in a combinatorial explosion can have the same net conversion. Thus, the number of net conversions is generally much smaller than the number of net conversions. Since net conversions cannot yet be independently computed from EFMs, the smaller set of net conversions needs to be deduced from the redundant set of EFMs.

## Discussion

This paper serves as an educational text on the determination and analysis of elementary flux modes and their associated net conversions. Net conversions play an important role in microbial ecology and metabolic engineering and can be found in classical textbooks [1, 2, 22]. They express the yields of the pathway products, e.g. ATP and catabolic byproducts, on the pathway substrate and allow for a bioenergetic evaluation of the performance of a pathway. Microbes can express a multitude of different net conversions, or mixtures thereof, each adapted to different conditions. Explaining the metabolic strategies exhibited by microbes in different environments remains a key objective of microbiology [7].

The methods for computing net conversions include a heuristic manual method, one based on linear algebra, and another that is entirely computational and depends on a mathematical definition of a metabolic pathway: an elementary flux mode [23, 11]. Perhaps the simplest definition of an EFM was offered by one of its discoverers, Stefan Schuster: ‘An elementary flux mode is a minimal set of enzymes that could operate at steady state, with all the irreversible reactions used in the appropriate direction’ [24]. Thus, an EFM network is a network of reactions that is minimal, indicating that: i. no reactions can be removed without violating the steady-state condition for all the internal intermediates, ii. none of the reactions are redundant, and iii. therefore it has a single independent flux.

EFMs have a number of appealing properties for applications [11, 12]. Firstly, the metabolic pathway with maximal yield, e.g. of a catabolic product or ATP on the energy source, is always an EFM [15]. For this reason, flux balance analysis, which computes maximal yield pathways, and EFMs are intimately related. Secondly, all feasible steady-state rate vectors of any metabolic network are always expressible as combinations (with positive weight coefficients; convex or conic combination) of the steady-state rate vectors of an EFM. Thus, all possible steady-state flux distributions of a network can be reconstructed from all its EFMs. Finally, since one EFM is associated with a single net conversion, the set of net conversions possible given a metabolic network arises from the combinations of the net conversions of its constituent EFMs.

One condition of an EFM is that all its reactions operate in their ‘allowed’ or ‘feasible’ (physiologically or thermodynamically) direction. Thus, irreversible reactions such as kinases are only allowed in their feasible direction in an EFM. (This ensures that all of the computed EFMs are in principle attainable by the cell.) We achieved this so far implicitly by assigning the import reaction of the energy source a rate value of 1, so of import instead of production. This implies that when none of the networks shown in Figures 3 and 4 had irreversible reactions, each of these networks would give to two EFMs one operating from the energy substrate to the catabolic product and the other operating in the opposite direction, from product to substrate. This illustrates also that the nullspace calculation which led to a single flux vector is not equivalent to an EFM computation. An EFM computation requires, in addition to steady-state flux balances, the identity of the irreversible reactions of the network. A network with more than 1 independent flux generally gives rise to more than one EFM (depending on which reactions are considered irreversible). And since each EFM is associated with a single net conversion, the steady-state flux vector corresponding to an EFM leads to a net conversion according to the calculation started at equation 1. The Python package EFMTool calculates EFMs. We use this package also in our pipeline to compute net conversions and their thermodynamic information.

We also presented a straightforward thermodynamic measure of a net conversion and its underlying pathway. Currently, in textbooks, we find only that the Δ*G*^*m′*^ or Δ*G*^0*′*^ is reported. In academic papers either the maximal values of the minimum driving force are mentioned or the free energy differences in the enzyme-cost minimum. However, when standard free energies are missing or when intracellular concentration bounds cannot be confidently estimated our measure is useful (it then also equals the MDF result in the absence of concetration bounds). This is the case in, for example, the TCA cycle, where the concentration of oxaloacetate was required to be below the lower bound to find an MDF where all Δ*G*’s were negative [20].

We hope that this paper motivates the continued study of the metabolic potential and adaptive strategies of microorganisms using rigorous and quantitative methods that ensure unambiguous analysis of metabolic stoichiometry and bioenergetics.

## Methods

### Python pipeline to compute EFMs, net conversions and their Gibbs free change

We present an intuitive Python pipeline as a Python notebook that determines for any metabolic network: i. its EFMs, ii. their net conversions, and iii. the Gibbs free energy change under standard conditions. The pipeline is based on existing tools such as cobrapy [25], EFMTool [13] and eQuilibrator [26]. It is available as a Python notebook on GitHub and requires minimal Python coding expertise. It can be run in a browser using Google Colab and does not require any Python installations. If actions are required by the user, there is a ‘TO DO’ box with a description of the required action. The notebook works for different metabolic networks and we provide several examples as supplemental information.

The following steps are performed in the Python notebook:

1. **Provide spreadsheet with pathway information**. This is the most labor-intensive step of the pipeline. A spreadsheet with two tabs, one for reactions and one for metabolites, is the input of the pipeline. The minimal information required for the reactions is a reaction ID, stoichiometry, and reversibility. The minimal information required for the metabolites is a metabolite ID and a KEGG [18] identifier (or elemental composition). The input file for EMP glycolysis with lactate production is given in Supplementary file S2. Inputs in the Python notebook are a list of allowed external metabolites. In the implementation in Python (Supplementary file S3), the convention of exchange reactions is used instead of removal of the external metabolites from the stoichiometric matrix is used for easy calculation of overall equations. An exchange reaction is a reaction where an (external) metabolite is consumed into ‘nothing’. The two approaches are completely equivalent.
2. **Check the elemental balance of the reactions**. In the pipeline, there are built-in functions that check whether all reactions in the input spreadsheet are elementally balanced. The results of these checks are presented in df_balance, which should be inspected. If imbalanced reactions are present, they should be corrected in the spreadsheet and the spreadsheet should be reloaded in the code.
3. **Build the COBRApy model and perform basic verification**. The metabolites and reactions are loaded into a COBRApy model. Then, some basic network verification is performed using flux balance analysis: production of all products from the substrate is optimized and ATP production from ADP and Pi, without the presence of substrate is optimized. If the model does not behave as expected, e.g. infinite ATP can be produced without input of substrate, the input spreadsheet should be adapted until the model behaves as expected. If there are problems with the network, analysis of blocked reactions can help identify the issues. Optionally, the model can be exported as sbml and/or json file.
4. **Enumerate the EFMs using EFMTool**. The Python implementation of EFMTool is used to enumerate the EFMs in the network. The input for EFMTool is the stoichiometric matrix of the network and an array of reversibility of the reactions, which can be extracted from the cobrapy model made in the last step. If the network consists of a single EFM, the output EFM is the full network. However, if there are redundant reactions, these are not part of the EFM. (In the example of lactate production from glucose, there is no net production nor consumption of protons. Therefore, the proton transport reaction and exchange reaction are not utilized in the EFM.) If there are multiple EFMs in the network, EFMTool outputs all EFMs in an array, where the rows are the reactions, and each column is an EFM. For practical reasons, the EFMs are normalized for a certain reaction, in this case, the uptake of glucose. The output of EFMTool is normalized in a manner that the smallest flux has value 1. For biological interpretation, normalization by the substrate uptake, or ATP production are sensible alternatives.
5. **Determine the net conversions of the computed EFMs**. The overall conversion of an EFM is calculated by considering the value of the flux through all exchange reactions. The flux values signify the yield on the metabolite that was normalized for.
6. **Determine the Gibbs free energy change of the overall reactions of the computed EFMs**. The Gibbs free energy change under standard conditions is calculated for the net conversion with eQuilibrator.

For each of the networks presented in this work, e.g. Figures 1-5, a supplementary Python notebook file, combined with an input spreadsheet, is available which performs these steps.

### NetConversionApp and KEGGBuilderApp

The apps and instructions how to install them are available at https://github.com/SystemsBioinformatics/net_conversion_gui. Tutorial videos for both apps are available via the GitHub link. The Supplementary files, i.e. the Python notebook files and spreadsheets,can be found at https://github.com/SystemsBioinformatics/EFM_example_scripts.

## Acknowledgements

The authors thank Gustavo Jeuken for help in constructing the applications and Timothy Páez Watson for proofreading the document. The figures were compiled in with BioRender.com. Maaike Remeijer and Frank Bruggeman received support from NWO-XL grant OCENW.XL21.XL21.007 ‘Taking Control of Metabolism in Microbial Cell Factories by Applying Noncanonical Redox Cofactors’. Christoff Odendaal received salary support from the European Union’s Horizon research and innovation programme under Grant Agreement No 10113540 (Nutritive).

## Notes

https://github.com/SystemsBioinformatics/net_conversion_gui

https://github.com/SystemsBioinformatics/EFM_example_scripts

### Competing Interest Statement

The authors have declared no competing interest.

