## Appendix for "Net conversion calculations of catabolic pathways"

### S1 Step by step determination of all steady state rate in figure 1

We start by assigning the rate of glucose uptake reaction (PTS), denoted by $v_{PTS}$, a value of 1 (mmol/(gram dry weight $\times$ hour)). For the concentration of $G6P$, denoted by $[G6P]$, to be constant (hence, for $d[G6P]/dt = 0$) it is required that $v_{GPI}$ also equals 1. The constancy of $[F6P]$ and $[F16BP]$ requires $v_{PFK} = v_{ALDO} = 1$. Aldolase splits fructose-1,6-bisphosphate, $F16BP$ – a six carbon atom and two phosphate containing molecule – into two molecules that are isomers ($G3P$ and $DHAP$), each containing 3 carbon atoms and a single phosphate group. The enzyme $TPI$ interconverts these isomers. The enzyme $GAPDH$ then processes $G3P$ further. The constancy of $[DHAP]$ requires that $v_{TPI} = 1$. The net synthesis rate of $G3P$ equals $v_{ALDO} + v_{TPI} = 2$ such that $v_{GAPDH} = 2$ to ensure constancy of $[G3P]$. Ensuring the constancy of $[BPG]$, $[PG3]$, and $[PG2]$ then requires that $v_{PGK} = v_{PGM} = v_{ENO} = 2$. This implies that the synthesis rate of $PEP$ equals 2. The PTS system for glucose uptake that we are considering has $PEP$ as substrate and runs at rate $v_{PTS} = 1$, which implies for the constancy of $[PEP]$ that the rate of enzyme PYK equals 1. Since the PTS system has $PYR$ as product, its net synthesis rate equals $v_{PTS} + v_{PYK} = 2$ and the rate of $LDH$ needs to equal 2 to ensure constancy of $[PYR]$. Now we have balanced all the internal metabolites except for $NADH$ and $NAD$. The concentration of $NADH$ is indeed constant, since its rate of synthesis by $GAPDH$ is balanced by its consumption rate by $LDH$; they are both equal to 2. Since in each $NADH$ producing (or consuming) reaction an equal amount of $NAD$ is consumed (and produced), also the constancy of $[NAD]$ is ensured.

### S2 Thermodynamics primer

Thermodynamics defines the Gibbs free energy of formation ($G$) as the energy in Joules $J$ needed to form 1 mole of a specific chemical compound from chemical elements under standard conditions (and at constant pressure and temperature). Accordingly, the change in the Gibbs free energy of a reaction (named 'r')

$$aA + bB \overset{r}{\rightleftharpoons} cC + dD,$$

with all the stoichiometric coefficients in small font, equals

$$\Delta G_r = cG_C + dG_D - aG_A - bG_B.$$

Where $G_i$ denotes the Gibbs free energy of formation of compound $i$, also commonly denoted as $\Delta_f G_i$.

The value of $\Delta G_R$ is negative when less energy was needed to form the products under standard conditions than to form the substrates under those same conditions and, hence, Gibbs free energy is lost ('dissipated'). Below we will see that this lost energy is not wasted, because it accelerates the rate of the reaction.

The Gibbs free energy of formation of a chemical compound, say $A$, can be calculated from

$$G_A = G_A^{0\prime} + RT\ln[A]$$

with $G_A^{0\prime}$ as the energy requirement in $J/mol$ to make 1 mole of $A$ from the reference of set of compounds under standard conditions, $R$ as the ideal gas constant (8.314 $J/K$), and $T$ as the temperature in $K$.

From these definitions it follows that the Gibbs free energy change of a reaction can be expressed in terms of the concentrations of the reactants,

$$\Delta G_r = \Delta G_r^{0\prime} + RT\ln\frac{[C]^c[D]^d}{[A]^a[B]^b},\quad \Delta G_r^{0\prime} = cG_C^{0\prime} + dG_D^{0\prime} - aG_A^{0\prime} - bG_B^{0\prime}.$$

This last equation directly relates to the rate equation of reaction, defined in kinetic theory. The rate $v_r$ of reaction $r$ equals the difference between its forward rate $v_r^+$ and backward rate $v_r^-$,

$$v_r = v_r^+\left(1-\frac{v_r^-}{v_r^+}\right) = \kappa f([\text{reactants}],\ [\text{effectors}])\left(1-\frac{[C]^c[D]^d}{[A]^a[B]^bK_{eq}}\right),\ K_{eq} = e^{-\frac{\Delta G^{0\prime}}{RT}}$$

with $\kappa$ as a function of kinetic constants that is different for mass-action and enzyme kinetics and $f([\text{reactants}],\ [\text{effectors}])$ as the saturation-with-substrate function. This function equals in the case of enzyme kinetics, for instance,

$$0 \le \frac{\frac{[A][B]}{K_AK_B}}{\left(1+\frac{[A]}{K_A}+\frac{[C]}{K_C}\right)\left(1+\frac{[B]}{K_B}+\frac{[D]}{K_D}\right)} \le 1,\ \text{for: } a=b=c=d=1,$$

while for mass action kinetics it equals $[A]^a[B]^b \ge 0$.

From the above, it follows that

$$v_r = \kappa f([\text{reactants}],\ [\text{effectors}])(1-e^{\frac{\Delta G_r}{RT}}).$$

This last equation indicates that

$$\Delta G_r < 0 \ \Leftrightarrow\ v_r > 0,$$

$$\Delta G_r = 0 \ \Leftrightarrow\ v_r = 0,$$

$$\Delta G_r > 0 \ \Leftrightarrow\ v_r < 0,$$

Thus, the sign of the Gibbs free energy change determines whether the reaction proceeds forwards or backwards. When $\Delta G_r$ is zero, the rate is zero and the reaction operates at thermodynamic equilibrium ($v_r^+ = v_r^- \neq 0$). This sign relation ensures that entropy is always produced since $dS/dt = -v_r\Delta G_r/T \geq 0$ with T as the absolute temperature, in agreement with the second law of thermodynamics.

Because $1-e^{\frac{\Delta G_r}{RT}}$ increases (approximately hyperbolically) with $-\Delta G_r$, more negative values of $\Delta G_r$ are expected to give rise to higher reaction rates; in particular because $f$([reactants], [effectors]) generally increases also with $-\Delta G_r$. Finally, because $1 - e^{\frac{\Delta G_r}{RT}}$ equals 0 at thermodynamic equilibrium and approximates 1 when $\Delta G_r \to -\infty$, it is often referred to as the displacement from thermodynamic equilibrium. Thus, a reaction proceeds faster when it is further displaced from thermodynamic equilibrium.

### S3 Ω measure

We start from the following information,

$$\{\{\Delta G_1, v_1\}, \{\Delta G_2, v_2\}, ..., \{\Delta G_i, v_i\}, ...\{\Delta G_{nR}, v_{n_R}\}\} \tag{S1}$$

and relationships,

$$\Delta G_{CAT} = \sum_{i=1}^{n_R} \frac{v_i}{v_{NC}} \Delta G_i = n_R \Big\langle \frac{v}{v_{NC}} \Delta G \Big\rangle = n_R \left( \Big\langle \frac{v}{v_{NC}} \Big\rangle \langle \Delta G \rangle + covar(v/v_{NC}, \Delta G) \right), \tag{S2}$$

with $\langle x \rangle = \frac{\sum_{i=1}^{n} x_i}{n}$ as the mean of the values $\{x_1, x_2, ..., x_i, ..., x_n\}$ and $covar(x, y)$ as the covariance between between $x$ and $y$ values. We used the covariance relation: $covar(xy) = \langle xy \rangle - \langle x \rangle \langle y \rangle$.

Next, we assume that all flux values $v_i$ are positive, which can always be achieved by reverting the direction of reactions), such that all $\Delta G_i$ are negative. Next, we derive,

$$\Omega = \frac{-\Delta G_{CAT}}{n_R \Big\langle \frac{v}{v_{NC}} \Big\rangle} = -\sum_{i=1}^{n_R} \omega_i \Delta G_i, \quad \omega_i = \frac{v_i}{\sum_{i=1}^{n_R} v_i}, \ \forall i: \ \omega_i \geq 0, \ \sum_{i=1}^{n_R} \omega_i = 1 \tag{S3}$$

with $\omega_i$ as the flux fraction of reaction $i$ such that $\Omega$ can be interpreted as a flux-weighted average.

This last expression leads to the following inequality,

$$\Omega \geq -\min(\Delta G_i), \tag{S4}$$

with $\min(\Delta G_i)$ as the minimum $\Delta G_i$ value. Equality holds when all $\Delta G_i$'s are identical. This is the result of a max min driving force (MDF) optimization in which none of the concentrations bounds are active, .i.e. the true maximal objective value can be attained.

From equation S2, we derive that

$$\Omega = \langle -\Delta G \rangle \left( 1 + \frac{covar(v/v_{NC}, -\Delta G)}{\langle v/v_{NC} \rangle \langle -\Delta G \rangle} \right), \tag{S5}$$

when $covar(v/v_{NC}, -\Delta G) \geq 0$ which is the expectation (since rates are higher when their $\Delta G$ is more negative),

$$\Omega \geq -\langle \Delta G \rangle \tag{S6}$$

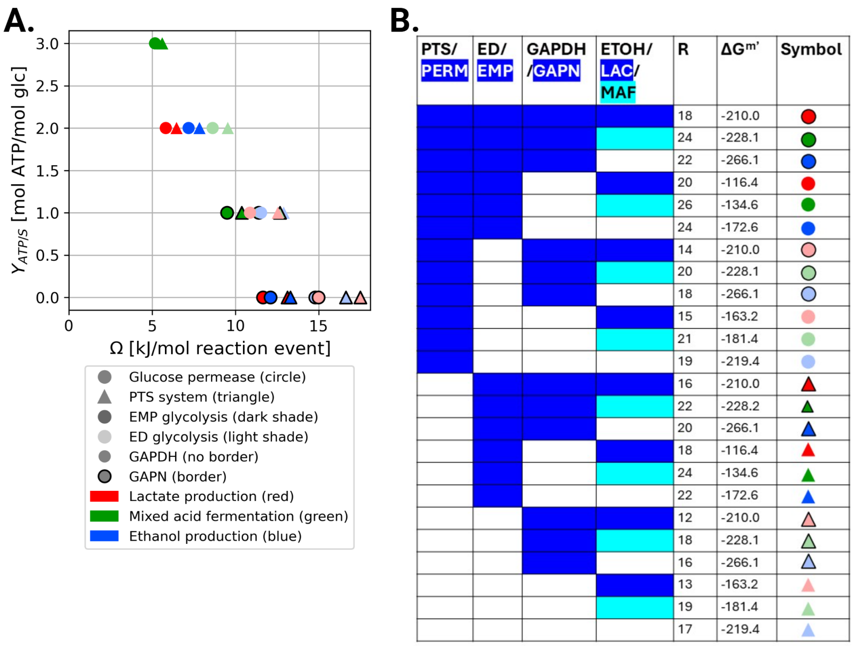


Figure S1: **Pathway ranking based on the Ω measure.** (A) ATP yield as a function of Ω for the 24 EFMs in the pan-glycolysis network. The shape, colour, shading and border of the datapoints indicates the combination of transporter, product spectrum, EMP/ED and GAPDH variant of each EFM. (B) Extended legend for figure A. R indicates the number of reaction steps that the $\Delta G^{m\prime}$ is divided by. The $\Delta G^{m\prime}$ is given in kJ/mol, calculated using the eQuilibrator API.

It follows that

$$-\langle \Delta G \rangle \geq -\min(\Delta G_i), \tag{S7}$$

which reduces to an equality when all $\Delta G_i$ are identical. Finally, we from equations S4, S6, and S7 that

$$\Omega \geq -\langle \Delta G \rangle \geq -\min(\Delta G_i). \tag{S8}$$

The usefulness of the Ω measure is that it allows us to rank pathways in terms of units of flux sustained per unit of enzyme investment *assuming that all enzymes have comparable kinetics*. As an example, for the pan-catabolic network shown in Figure 5, see the Ω in Figure S1. The catabolic pathway with the highest Ω therefore has the highest flux/enzyme investment, assuming identical kinetics ($k_{cat}$ and $K_M$ values). We do not know the kinetics of most enzymes, so we cannot directly calculate the *true* enzyme investment to sustain a unit of flux. If one pathway contains catalytically efficient enzymes (higher $k_{cat}$)

than another, for example, the former will require a lower enzyme investment to sustain the same flux. However, in the absence of accurate and detailed enzyme kinetics, the $\Omega$ measure provides a way to estimate relative merit of the pathway, assuming that there are no large differences in enzyme kinetics. For this, we need (and have): all reaction stoichiometries, $\Delta G_i^{m\prime}$, $n_R$, and $v_i/v_{NC}$ such that we can calculate the net conversion, $\Delta G_{cat}$, $\langle \Delta G_i^{m\prime} \rangle$, $\langle v/v_{NC} \rangle$ and, finally, $\Omega$.

### S4 $\Omega$ is equivalent to MDF without concentration bounds

Above, $\Omega$ was introduced. Here, we show that it is equal to the solution of max-min driving force (MDF) analysis, when no concentration bounds are applied, if the pathway analyzed is a ray (an EFM that is not a cycle without net conversion). The MDF is calculated by the following linear program [1]:

$$
\begin{aligned}
\max_{B,\,\mathbf{y}} \quad & B \\
\text{s.t.} \quad & \Delta_r G_i^{\circ\prime} + \Delta_r G_i^{\text{bnd}} + RT \sum_{j \in \mathcal{M}_{\text{int}}} S_{ij}\, y_j \;\leq\; -B \qquad \forall\, i \in \mathcal{R} \\
& \ln x_j^{\text{lb}} \;\leq\; y_j \;\leq\; \ln x_j^{\text{ub}} \qquad \forall\, j \in \mathcal{M}_{\text{int}} \\
& B \in \mathbb{R}, \quad y_j \in \mathbb{R}
\end{aligned}
\tag{S9}
$$

Which contains the following symbols:

- $B$ — the minimum driving force to be maximised (kJ mol$^{-1}$)
- $y_j = \ln[x_j]$ — log-concentration of internal metabolite $j$ (dimensionless)
- $\mathcal{R}$ — set of reactions; $\mathcal{M}_{\text{int}}$ — set of internal metabolites
- $S_{ij}$ — stoichiometric coefficient of metabolite $j$ in reaction $i$
- $\Delta_r G_i^{\circ\prime}$ — standard transformed Gibbs energy at pH 7, $I = 0.1\,\text{M}$ (kJ mol$^{-1}$)
- $\Delta_r G_i^{\text{bnd}} = RT \sum_{j \in \mathcal{M}_{\text{bnd}}} S_{ij} \ln x_j^{\text{fix}}$ — correction from fixed boundary metabolite concentrations
- $\Delta_r G_{i,\,\text{eff}}^{\circ\prime} = \Delta_r G_i^{\circ\prime} + \Delta_r G_i^{\text{bnd}}$ — effective standard Gibbs energy after absorbing boundary concentrations
- $x_j^{\text{lb}}$, $x_j^{\text{ub}}$ — lower and upper concentration bounds on internal metabolite $j$
- $R$ — universal gas constant (kJ mol$^{-1}$ K$^{-1}$); $T$ — absolute temperature (K)

B, or the MDF value, is the value of the driving force of the reaction in the pathway that has the lowest driving force ($-\Delta G$). Multiplying the constraint indicating all reactions must have a driving force equal to or lower than B with the flux vector, gives the following:

$$\sum_{i\in\mathcal{R}} v_i\, \Delta_r G^{\circ\prime}_{i,\text{eff}} + RT \sum_{j\in\mathcal{M}_{\text{int}}} y_j \underbrace{\sum_{i\in\mathcal{R}} v_i S_{ij}}_{=0} \;\leq\; -B \sum_{i\in\mathcal{R}} v_i \tag{S10}$$

Because an EFM is by definition mass balanced, $\sum_{i\in\mathcal{R}} v_i S_{ij} = 0$ for all internal metabolites $j$. Note that boundary concentrations (the concentrations of the metabolites in the overall conversion) and concentration bounds (the minimal and maximal concentrations assumed for an internal metabolite) are two different concepts. The left-hand side of this equation can be expanded as follows:

$$\sum_{i\in\mathcal{R}} v_i\, \Delta_r G^{\circ\prime}_{i,\text{eff}} \;=\; \underbrace{\sum_{i\in\mathcal{R}} v_i\, \Delta_r G^{\circ\prime}_i}_{\Delta G^{\circ\prime}_{\text{CAT}}} + RT \sum_{j\in\mathcal{M}_{\text{bnd}}} \ln x^{\text{fix}}_j \underbrace{\sum_{i\in\mathcal{R}} v_i S_{ij}}_{\sigma_j} \;=\; \Delta G_{\text{CAT}} \tag{S11}$$

where $\sigma_j = \sum_i v_i S_{ij}$ is the net stoichiometric coefficient of boundary metabolite $j$ in the overall reaction, and $\Delta G_{\text{CAT}}$ is the Gibbs energy of the net conversion at the chosen boundary concentrations. Thus:

$$\Delta G_{CAT} \leq\; -B \sum_{i\in\mathcal{R}} v_i \tag{S12}$$

Assuming all $v_i \geq 0$, which is trivially achieved by splitting reversible reactions into a forward and reverse component, the following relationship is obtained:

$$B \;\leq\; \frac{-\Delta G_{\text{CAT}}}{\sum_{i\in\mathcal{R}} v_i} \;=\; \Omega \tag{S13}$$

If the internal metabolite concentrations are unconstrained, the $y_j$ can always be chosen in such a way that $B^* = \Omega$. This equality holds for any choice of boundary concentrations.

We verified this equality for EMP glycolysis with the production of lactate (Figure S2) and the TCA cycle (Figure S3). For both pathways, the $\Omega$ and MDF were calculated for the case where all boundary concentrations were 1 mM, and for a more physiological case, for which the exact concentrations are described in the figure legends. The $\Delta G^{m\prime}$ values were obtained using the eQuilibrator API [2]. In all cases, indeed the MDF without concentration bounds was equal to $\Omega$.

In the case of EMP glycolysis, the MDF with and without concentration bounds are close in value (Figure S2a), whereas the value of the MDF with concentration bounds of the TCA cycle is negative (i.e. not all reactions can have a

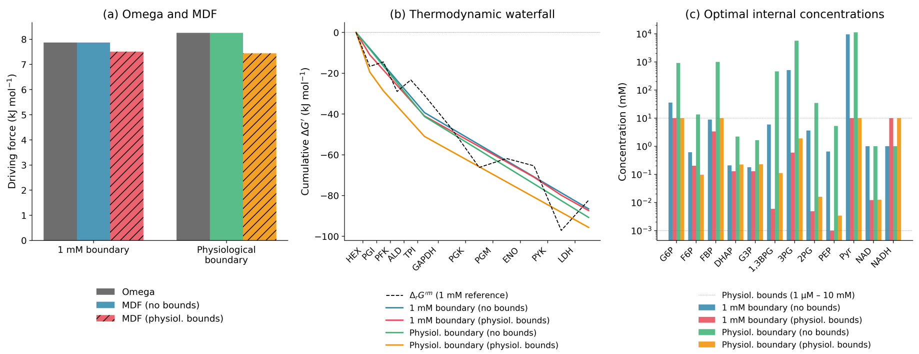


Figure S2: **Max-min driving force and Ω analysis of EMP glycolysis.** (a). The Ω, MDF without bounds and MDF with bounds for the situation where all boundary concentrations are 1 mM and for physiological boundary concentrations. Physiological boundary concentrations are 5 mM glucose, 1 mM lactate, 3 mM ATP, 0.5 mM ADP and10 mM phosphate. (b). Cumulative $\Delta G$ of each reaction. In the case of MDF without concentration bounds, the $\Delta G$ is equally distributed. (c). The optimal concentrations for all MDF simulations. Note that the concentrations are not unique, there may be multiple sets of concentrations that give rise to the same MDF value.

negative $\Delta G$ under these conditions), while Ω is positive. This was also reported in Noor et al. (2014) [1], which they corrected by lowering the concentration of oxaloacetate. In the MDF without bounds, the oxaloacetate concentration is low. The exact concentration values (Figures S2c and S3c) are not unique, there may be multiple concentration values that lead to the same optimal B. When there are concentration bounds, this is likely more restricted and fewer alternative optima exist. For the TCA cycle calculation, more intracellular concentrations were required as input (NAD(H), FAD($H_2$), CoA, acetyl-CoA), as it is not a pathway that converts an extracellular compound into another extracellular compound. This brings more uncertainty, both in the MDF and Ω analysis.

In this analysis, transporters were omitted for simplicity, but the results are equivalent if a transporter is included in the pathway. The boundary concentrations are in that case the extracellular metabolite concentrations instead of the intracellular ones.

[2] Moritz E Beber, Mattia G Gollub, Dana Mozaffari, Kevin M Shebek, Avi I

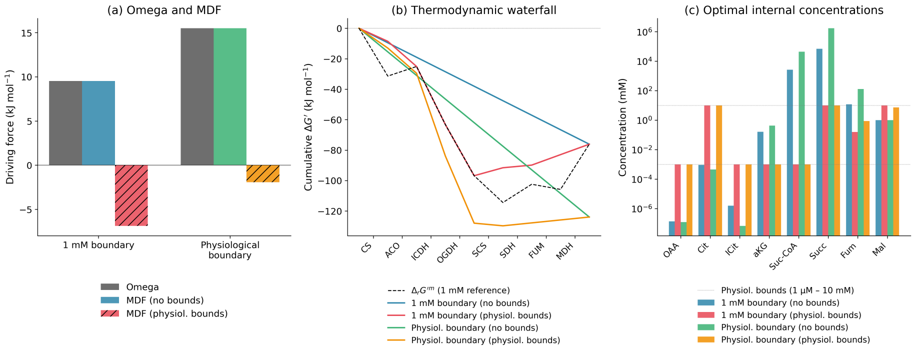


Figure S3: **Max-min driving force and Ω analysis of the TCA cycle.** (a). The Ω, MDF without bounds and MDF with bounds for the situation where all boundary concentrations are 1 mM and for physiological boundary concentrations. Physiological boundary concentrations are 0.6 mM acetyl-CoA, 0.1 mM CoA, 1.45 $\cdot 10^{-2}$ mM $CO_2$ (atmospheric, 445 ppm), 3 mM ATP, 0.5 mM ADP, 10 mM phosphate, 10 mM NAD, 1 mM NADH, 0.5 mM FAD and 0.1 mM $FADH_2$. (b). Cumulative $\Delta G$ of each reaction. In the case of MDF without concentration bounds, the $\Delta G$ is equally distributed. (c). The optimal concentrations for all MDF simulations. Note that the concentrations are not unique, there may be multiple sets of concentrations that give rise to the same MDF value.

Flamholz, Ron Milo, and Elad Noor. equilibrator 3.0: a database solution for thermodynamic constant estimation. *Nucleic acids research*, 50(D1): D603–D609, 2022.
